# Genomic positional conservation identifies topological anchor point (tap)RNAs linked to developmental loci

**DOI:** 10.1101/051052

**Authors:** Paulo P. Amaral, Tommaso Leonardi, Namshik Han, Emmanuelle Viré, Dennis K. Gascoigne, Raúl Arias-Carrasco, Magdalena Büscher, Anda Zhang, Stefano Pluchino, Vinicius Maracaja-Coutinho, Helder I. Nakaya, Martin Hemberg, Ramin Shiekhattar, Anton J. Enright, Tony Kouzarides

## Abstract

The mammalian genome is transcribed into large numbers of long noncoding RNAs (lncRNAs), but the definition of functional lncRNA groups has proven difficult, partly due to their low sequence conservation and lack of identified shared properties. Here we consider positional conservation across mammalian genomes as an indicator of functional commonality. We identify 665 conserved lncRNA promoters in mouse and human genomes that are preserved in genomic position relative to orthologous coding genes. The identified ‘positionally conserved’ lncRNA genes are primarily associated with developmental transcription factor loci with which they are co-expressed in a tissue-specific manner. Strikingly, over half of all positionally conserved RNAs in this set are linked to distinct chromatin organization structures, overlapping the binding sites for the CTCF chromatin organizer and located at chromatin loop anchor points and borders of topologically associating domains (TADs). These ***t***opological ***a***nchor ***p***oint (tap)RNAs possess conserved sequence domains that are enriched in potential recognition motifs for Zinc Finger proteins. Characterization of these noncoding RNAs and their associated coding genes shows that they are functionally connected: they regulate each other’s expression and influence the metastatic phenotype of cancer cells *in vitro* in a similar fashion. Thus, interrogation of positionally conserved lncRNAs identifies a new subset of tapRNAs with shared functional properties. These results provide a large dataset of lncRNAs that conform to the “extended gene” model, in which conserved developmental genes are genomically and functionally linked to regulatory lncRNA loci across mammalian evolution.

## 1 Introduction

Long noncoding RNAs (lncRNAs) comprise the main transcriptional output of the mammalian genome, with recent surveys cataloguing approximately 60,000 lncRNA genes in humans compared to some 21,000 protein-coding loci (Iyer et al., 2015). While many of these lncRNAs are transcribed by RNA Polymerase II and are predominantly spliced and polyadenylated in a similar manner to protein-coding genes, there are currently no sequence or structural features that are predictive of their biological functions.

The functions of only a small fraction of lncRNAs have been experimentally tested, predominantly on an *ad hoc* basis, revealing that they can exert different roles to regulate genome function and gene expression at different levels (Amaral and Mattick, 2008; Flynn and Chang, 2014). From a mechanistic point of view, lncRNAs can act both post-transcriptionally and at the level of chromatin structure and transcription, where they can affect the genes in the immediate genomic vicinity (*in cis*) and/or in other genomic locations (*in trans*) to repress or promote expression. Well-studied repressors include lncRNAs associated with imprinted loci such as *AIRN* (*Antisense IGF2R ncRNA*) and *KCNQ1OT1* (*KCNQ1 opposite strand/antisense transcript 1*), which promote the silencing of genomically associated genes in an allele-specific manner (Kanduri, 2015). Examples of activator RNAs include the lncRNAs *HOTTIP* (HOXA transcript at the distal tip) and transcripts with “enhancer-like function” such as ncRNA-a, which promote the transcription of neighboring genes (Guil and Esteller, 2012; Lai et al., 2013; Wang et al., 2011).

In contrast to coding genes, most lncRNAs do not exhibit high levels of primary sequence conservation across species (Carninci et al., 2005; Iyer et al., 2015). In fact, the increasing catalog of these transcripts such as *AIRN* and *XIST* (*X inactive-specific transcript*), which have been extensively molecularly characterized, indicate that they evolve under different functional constraints and exhibit high evolutionary plasticity (Pang et al., 2006). There are other indications as to what some of these differing constraints may be, including the early observation that lncRNAs can have promoters that exhibit higher conservation and that extends to longer sequence stretches than observed for promoters of coding genes (Carninci et al., 2005).

Using cDNA and EST genomic mapping, it has been shown that some lncRNAs have highly conserved promoters that are genomically localized in syntenic regions, which are consistently associated with orthologous genes in different species and these often have developmental functions (Dinger et al., 2008; Engstrom et al., 2006). For example, the lncRNA *SOX2OT* (*SOX2 Overlapping Transcript*) has alternative syntenic transcripts associated with highly conserved promoters in all vertebrate groups, with similar expression patterns, particularly in the central nervous system (Amaral et al., 2009). Importantly, other characterized lncRNAs that fall in the same category, such as *HOTTIP, WT1-AS* and *Evf2/Dlx6os* (*Dlx6 opposite strand transcript*) were demonstrated to participate in the regulation of the associated genes (Dallosso et al., 2007; Feng et al., 2006; Wang et al., 2011), suggesting a conserved functional connection between lncRNAs and neighboring genes. More recently, transcriptomic studies have expanded these observations and shown that synteny is observed for hundreds of lncRNAs across the genomes of amniotes and beyond (Hezroni et al., 2015).

Here, we systematically characterize genomic positional conservation of lncRNAs in mammals and investigate whether this feature is indicative of their biological roles. We identify 1700 positionally conserved lncRNAs, transcribed from 665 conserved syntenic promoters and find that they are predominantly associated with developmental genes, with which they are co-expressed in a conserved tissue-specific manner. Characterization of these RNAs shows that they can positively regulate their associated gene, can affect cancer-linked phenotypes in a similar fashion to the associated gene, have conserved sequence motifs and are strongly associated with chromatin loops, which contact enhancer regulatory sequences. This analysis provides a rich resource for the in-depth characterization of the ill-defined family of lncRNAs and their relationship to the structure and function of chromatin topological organization.

## 2 Results

### 2.1 Identification of positionally conserved RNAs in human and mouse

There is a paucity of information regarding the organization of lncRNAs into groups with common functionality. Here we considered that the conserved position of lncRNAs relative to coding genes may reflect a basis for identifying common properties and for functional indexing. We therefore set out to identify spliced lncRNAs that are positionally conserved in mammalian genomes. We compiled a comprehensive catalog of human and mouse transcripts based on 1) Gencode annotation (Harrow et al., 2012); 2) human and mouse RNA-sequencing (RNA-Seq) from six matched tissues (brain, cerebellum, heart, kidney, liver and testis) (Brawand et al., 2011); and 3) RNA-Seq data from four similar human and mouse cell lines (embryonic stem (ES), leukemia, lymphoblast and muscle cells) produced by the ENCODE project (**Supplementary Table S1**) (The ENCODE Project Consortium, 2012; Djebali et al., 2012). In total, we processed 80 RNA-Seq datasets and successfully mapped 2.6 billion reads.

Our analysis pipeline is designed for the comprehensive identification of human and mouse transcripts from both Gencode and the RNA-Seq data, with evidence of splicing, no overlap with coding exons in the same transcriptional orientation and no significant coding potential (**Figure 1A**). Promoter sequences of human lncRNAs were then aligned to the mouse genome to identify syntenic lncRNAs (see Supplementary Methods), since it is known that the promoter of lncRNAs can be conserved, even though the RNAs often show little sequence conservation (Carninci et al., 2005). Syntenic lncRNAs were defined as positionally conserved if their promoters were genomically associated with orthologous genes and produced spliced lncRNAs in the same relative transcriptional orientation (either sense or antisense relative to the coding gene) in both mouse and human (see Supplementary Methods).

**Figure 1:**
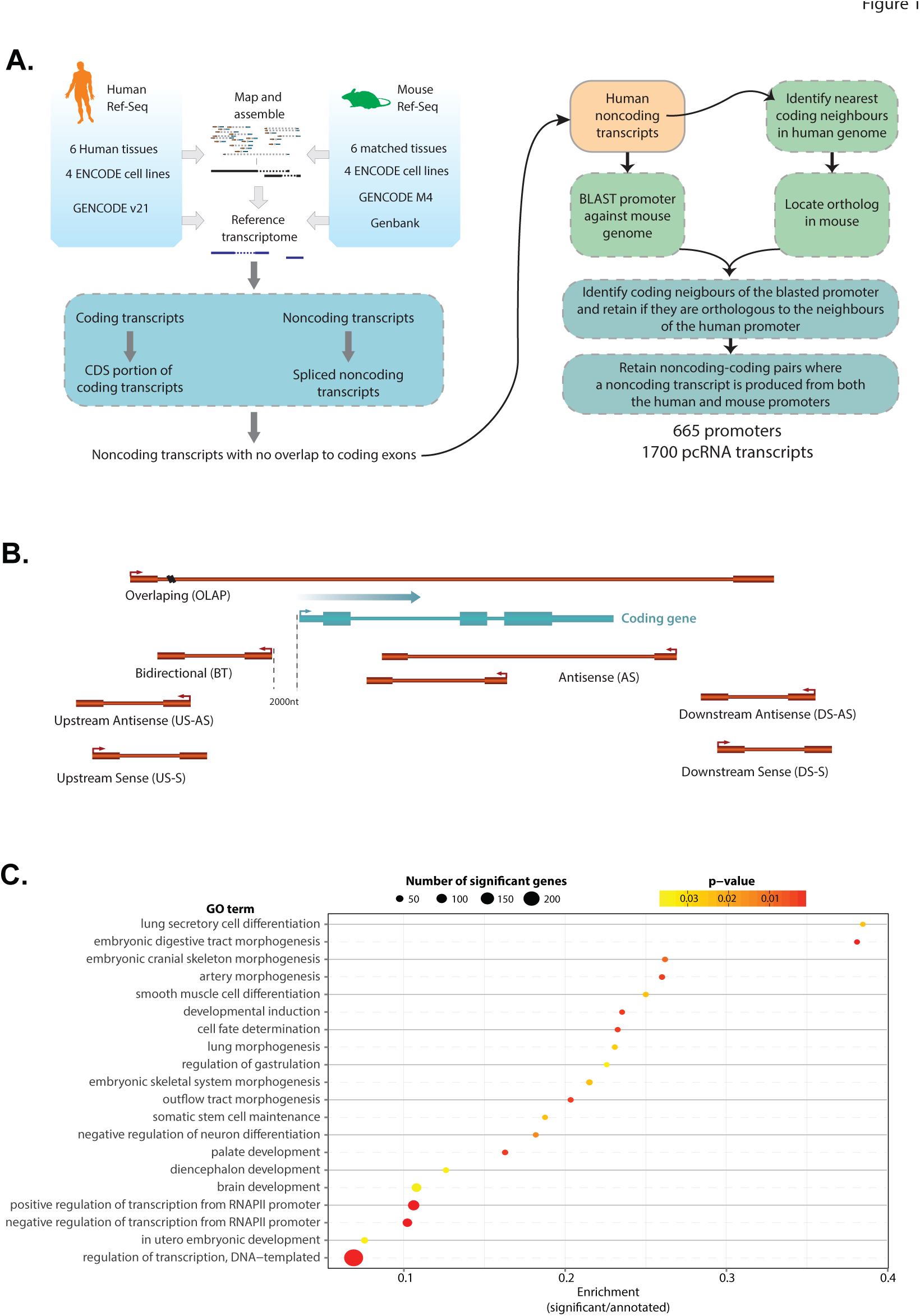
***A**: Workflow used for the identification of pcRNAs. **B**: Schematic diagram showing the possible orientations of a pcRNA (red) relative to a coding gene (blue). **C:** GO enrichment analysis of pcRNA-associated coding genes. The x-axis shows the enrichment score, calculated as the number of pcRNA-associated genes in a given GO category divided by the total number of genes in the category. The size of the points indicates the absolute number of pcRNA-associated genes in the given GO category. The color-coding indicates the adjusted p-value*.

The resulting set of positionally conserved lncRNAs (pcRNAs) comprises 1700 transcripts, including splicing isoforms, associated with 665 unique conserved promoters and a total of 626 orthologous coding genes (**Supplementary Table S2**). The majority of pcRNAs (82%, 1401/1700 transcripts, transcribed from 595 independent promoters) were Gencode annotated human transcripts, while 299 (18%) represented novel transcripts assembled from the RNA-Seq data. We also found that a small number of pcRNAs (138, transcribed from 32 independent promoters) overlapped syntenic microRNA (miRNA) loci, likely representing primary miRNA transcripts.

We classified pcRNAs into the following seven categories based on their genomic orientations relative to the associated coding genes (**Figure 1B**): Antisense (AS, direct overlap with the coding gene but transcribed from the opposite strand), Bidirectional (BT, transcribed in the opposite orientation with the transcription start site (TSS) within 2kb upstream the TSS of the coding gene), Overlapping (OLAP, partially overlapping the coding locus in the same orientation, with no overlap to coding exons and less than 50% overlap with untranslated regions), and transcripts Upstream (US) or Downstream (DS) of the coding genes in either sense (S) or antisense (AS) orientation.

The predominant category is Bidirectional transcripts (42% of all pcRNAs), followed by Antisense (18%), with all other categories similarly represented (between 5% and 9%) (**Supplementary Figure S1A**). On average, pcRNAs are 1.3kb long and have 3-4 exons (mean 3.6 exons per pcRNA), with most having 2 exons (**Supplementary Figure S1B,C**). In terms of genome distribution, most pcRNAs are proximal to coding genes. However, the distance between the pcRNA promoters and the TSS of the associated coding genes varies according to the positional category of the pcRNAs (**Supplementary Figure S1D**). Approximately 70% of pcRNAs are within 12kb of the associated genes. While BT, AS and OLAP promoter positions are, as expected, closest (median TSS to TSS distances of 215bp, 520bp and 35bp, respectively), promoters of upstream and downstream transcripts are usually much more distal (median TSS to TSS distances of 154kb (DS-S), 100kb (DS-AS), 44kb (US-AS), and 49kb (US-S)). Despite being less conserved then their associated coding genes, we found that human pcRNAs have on average 31% sequence identity with their mouse counterparts (**Supplementary Figure S1E**).

### 2.2 Positionally conserved RNA genes are associated with genes encoding developmental transcription factors

Gene Ontology (GO) analysis of the 626 coding genes with which pcRNAs are associated showed a very strong enrichment for genes with roles in *Regulation of transcription from RNA polymerase II promoter* (GO:0045944 and GO:0000122, adjusted p-values=1.2 × 10^−15^ and 8.5 × 10^−10^ respectively, **Figure 1C** and **Supplementary Table S3**). These are transcription factors involved in *Cell fate determination* (GO:0001709, adjusted p-value=0.00265) and *Developmental induction* (GO:0031128, adjusted p-value=0.00265), and central in the determination of a variety of specific developmental systems and embryonic stages, such as gastrulation regulation, stem cell maintenance and organ morphogenesis (**Supplementary Table S3**). Notably, many of these genes belong to major gene families containing regulators of lineage specification, such as *SOX* genes (including *SOX1*, *2, 4, 9* and *21*); *FOX* genes (*FOXA2*, *D3*, *E3*, *F1*, *I* and *P4*); *HOX* genes (e.g. *HOXA1*, *A2*, *A3*, *A11*, *A13, B3, C5* and *D8*) and other homeodomain genes, as well as several nuclear receptors, such as *NR2E1*, *NR2F1* and *NR2F2* (**Supplementary Table S3**).

Coding genes associated with pcRNAs showed an over-representation for developmental genes irrespective of their relative orientation to the associated pcRNA. We also found that certain categories had specific enrichment relative to all pcRNA-associated coding genes (**Supplementary Table S4**). For example, genes associated with bidirectional and overlapping pcRNAs were found to be enriched for signal transduction and signaling pathways (such as *IGF2*, *TGFB2* and *PIK3R5*) (**Supplementary Table S4**).

### 2.3 Positionally conserved RNAs have similar tissue specific expression patterns in mouse and human

To quantify the expression of pcRNAs and their associated protein-coding genes across the tissues and cell types, we used publicly available RNA-Seq data. As previously reported by us and others for lncRNAs (Cabili et al., 2011; Derrien et al., 2012; Dinger et al., 2008; Ravasi et al., 2006), pcRNAs have low average levels of expression when considered across all tissues (**Supplementary Figure S2A**), and their expression profiles often peak in one or a few specific tissues and/or cell lines (**Figure 2A**, **Supplementary Figure S2B,C**).

**Figure 2:**
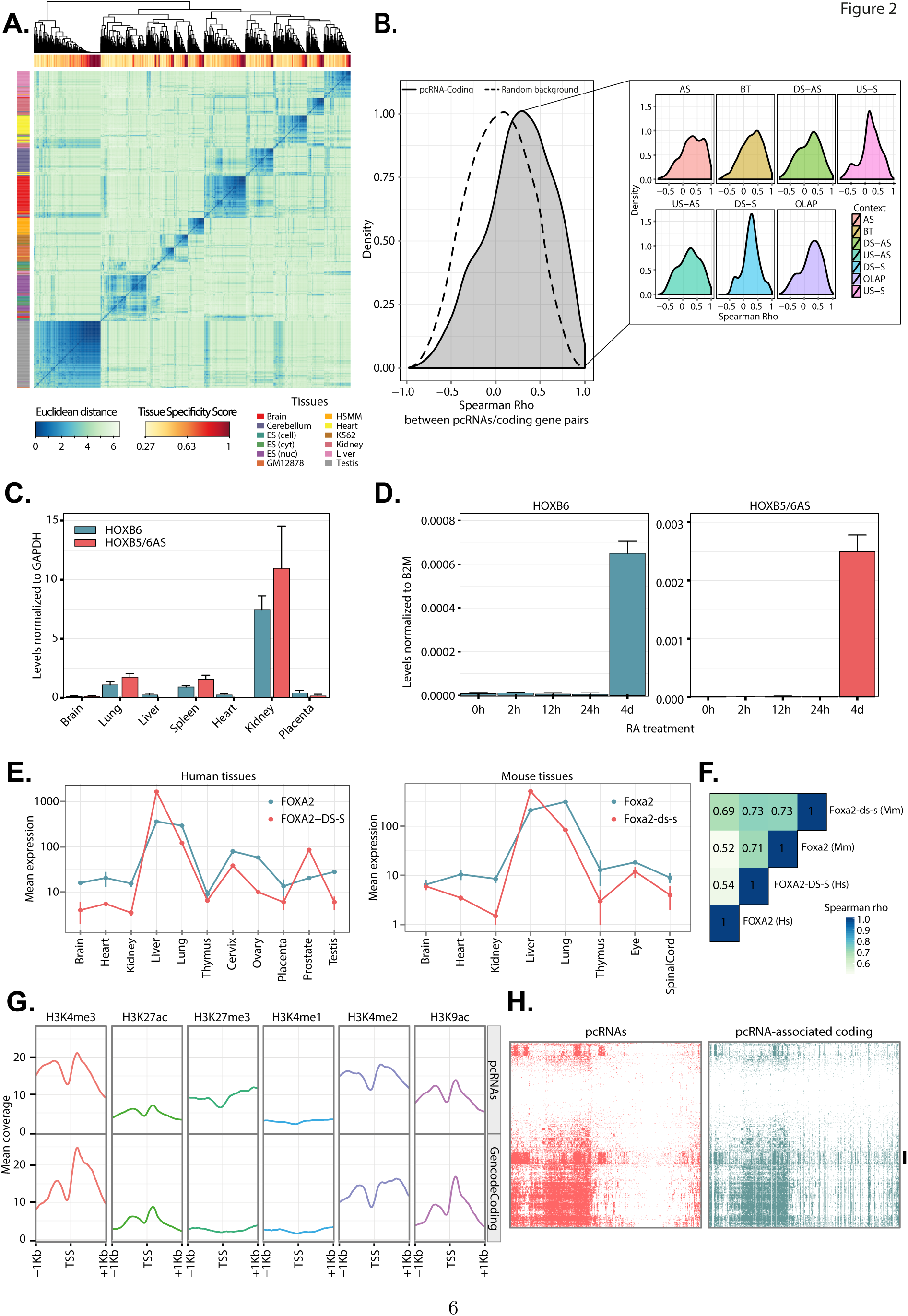
***A:** Heatmap showing the Euclidean distance between the expression profiles of human pcRNAs. The horizontal side bar reports the tissue specificity score of pcRNAs. The vertical sidebar reports the tissues in which each pcRNA has maximal expression. **B:** Density distribution of the Spearman’s correlation coefficients between pcRNAs and corresponding coding genes in human tissues and cell lines (mean Spearman’s rho 0.25, permutation test p-value <10^−6^). The dotted line shows the background distribution of all pairwise Spearman’s correlations between pcRNAs and pcRNA-associated coding genes. Inset: Distributions of the Spearman’s correlation coefficients dived by the positional category of the pcRNA. **C:** Real time PCR data showing the expression of* HOXB6 *(blue) and* HOXB5/6AS *(red) in a panel of 7 human somatic tissues. The data is expressed relative to the expression of* GAPDH*; the error bars indicate the standard error of the mean (SEM) across 3 technical replicate experiments. **D:** Real time PCR data showing the expression of* HOXB6 *(left) and* HOXB5/6AS *(right) over 5 time-points of NT2 cells differentiation with retinoic acid (RA). The data is expressed relative to the expression of* B2M*; the error bars indicate the standard error of the mean (SEM) across 3 replicate experiments. **E:** Nanostring expression profiles of* FOXA2 *and* FOXA-DS-S *across human (left) and mouse (right) tissues. The points indicate the mean value of two technical replicates, while the vertical bars report the value of each replicate. **F:** Heatmap showing Spearman’s correlation coefficients between the expression of human and mouse FOXA2 and FOXA-DS-S transcripts. **G:** Histone modification profiles of pcRNA promoters (top) and promoters of 1000 random Gencode coding genes (bottom) based on ChIP-Seq data by the ENCODE project on H1-hESCs. The lines represent the mean ChIP-Seq coverage. **H:** Transcription factor binding patterns in promoters of pcRNAs (left) and their associated coding genes (right). The heatmaps present the distribution of experimentally validated TF-binding sites from ENCODE 2,216 ChIP-seq experiments (y-axis), showing strong co-relations between the promoters of pcRNAs (x-axis) and their corresponding coding genes. The black bar indicates the binding pattern of the CCCTC-binding factor (CTCF)*.

We also found that pcRNAs exhibit significantly conserved expression patterns across mouse and human tissues (mean Spearman’s correlation 0.26, p-value<10^−6^, **Supplementary Figure S2D**) and very specific tissue expression signatures, with testis and brain/cerebellum containing the largest complements of tissue-specific pcRNAs (**Figure 2A**, **Supplementary Figure S2E**). Indeed, we identified numerous expression clusters of pcRNAs with highly correlated expression profiles and maximum expression in the same tissue, suggesting that pcRNAs may have conserved roles in tissue identity and cell type specification in mouse and human. Among the gene categories associated with tissue-specific pcRNAs expressed in any given tissue, we observed enrichment for regulatory genes involved in developmental and differentiation processes relevant to the particular tissue, such as neural differentiation genes in brain and endoderm developmental genes in liver (**Supplementary Figure S3A-D**).

### 2.4 Positionally conserved RNAs and genomically associated coding genes are co-expressed and co-induced

Expression of pcRNAs and associated genes showed a significant positive correlation when we analyzed RNA-Seq profiles across tissues (mean Spearman rho 0.25, p-value<10^−6^; **Figure 2B**), and in the majority of cases this correlation does not appear to be dependent on the proximity of the pcRNA to its coding gene (**Supplementary Figure S4A**). We also observed that pcRNAs have significantly higher tissue specificity than their associated coding genes (**Supplementary Figure S4B**). This result was also confirmed after taking into account the different expression levels of pcRNAs and coding genes, indicating that the increased tissue specificity of pcRNAs is not due to their lower expression level (see Supplementary Methods and **Supplementary Figures S4C,D**). These data suggest that pcRNAs and their corresponding coding genes are often co-regulated in mouse and human. Indeed, expression analysis of different pcRNAs in human NT2 (NTERA/D1) teratocarcinoma cells upon differentiation with all-trans retinoic acid – a widely-used system for regulation of Hox genes and ncRNAs (Sessa et al., 2007) - shows that they have regulated expression and are co-regulated with the associated coding genes, including homeobox genes, *NR2F1* and *TBX2* (**Figure 2C,D**, **Supplementary Figure S5**). For instance, *HOXB6* and *HOXB5/6-AS* transcripts are expressed in the same tissues and both can be induced when we differentiate NT2 cells (**Figure 2C,D**), similarly to the temporal co-induction seen with other coding/non-coding pairs (**Supplementary Figure S5**) and reminiscent of the co-regulation that we previously observed in differentiating mouse embryonic stem cells (Dinger et al., 2008).

To validate these results and extend to a larger and more diverse set of human and mouse tissues and cell types, we used the Nanostring expression assay, which provides a versatile platform for parallel single molecule detection of targeted RNAs with high sensitivity and specificity (Geiss et al., 2008). We designed a custom codeset to probe approximately 50 human and mouse manually selected pcRNAs and associated orthologous protein-coding genes, across an RNA panel of six matched human and mouse tissues. We also sampled additional tissues and cells such as mouse eye, spinal cord and embryonic stages; pluripotent cell lines from both species at various time-points of differentiation with retinoic acid, and a panel of 18 human cancer cell lines (**Supplementary Table S5**).

Using this approach we confirmed common co-expression of pcRNAs and associated genes (**Supplementary Figure S6A,B**), as well as conservation of expression between mouse and humans pcRNAs across different tissues (**Supplementary Figure S6C**). For example, *FOXA2* and its associated pcRNA *FOXA2-DS-S* exhibit very similar expression profiles across tissues both in human and mouse (**Figure 2E**), with Spearman correlation coefficients in the range 0.52-0.73 (**Figure 2F**). We observed similar results for *HNF1A* and its pcRNA *HNF1A-BT1/2* (**Supplementary Figure S6D-G**). Additionally, we found that pcRNAs often form clusters of co-expression with functionally related tissue-specific regulatory genes (**Supplementary Figure S7A,B**). For instance, two connected clusters are comprised of transcription factors that are master regulators of endoderm - in particular liver cell differentiation (*HNF1A*, *FOXA2*/*HNF3B* and *HNF4A*) (Costa et al., 2003; Odom et al., 2006) – and associated pcRNAs. These data suggest that pcRNAs share regulatory elements and/or have a role in the regulation of their neighboring protein-coding genes.

To investigate the principles of pcRNA regulation, we firstly inspected chromatin modification profiles around their TSS. Using the ENCODE genome-wide datasets for different human cell lines (ENCODE tier 1 lines: GM12878, H1-hESC, HSMM and K562) (The ENCODE Project Consortium, 2012; Hoffman et al., 2013), we identified a clear enrichment in H3K4 di-and tri-methylation (H3K4me2/3) and H3K9 and H3K27 acetylation (H3K9ac, H3K27ac) (**Figure 2G** and **Supplementary Figure S8**), as well as H3K27 tri-methylation (H3K27me3) and bivalent marks specifically for pcRNA promoters in ES cells (**Supplementary Figure S9**). At the same time, we observed only low levels of H3K4me1 in all the cell types analyzed. These profiles are indicative of RNA Polymerase II promoters, are similar to those observed for protein-coding genes (**Supplementary Figure S8**), and are not indicative of enhancers, suggesting that the pcRNA transcripts are not reflective of typical enhancer RNAs (eRNAs) produced from enhancer regions. Interrogation of the FANTOM5 Consortium database (Andersson et al., 2014), containing comprehensive annotations of over 40,000 enhancer regions, identified the promoters of only three pcRNAs as enhancers (associated with *GATA2*, *HES1* and *KLF4*) (data not shown).

### 2.5 Identification of topological anchor point (tap)RNAs

To obtain insights into the regulation of pcRNA and associated genes, we analyzed their promoters for transcription factor (TF) binding profiles. Using TF ChIP-seq peaks from the ENCODE project (The ENCODE Project Consortium, 2012; Wang et al., 2012), we observed a highly concordant pattern of occupancy in the promoters of pcRNAs and associated genes (Pearson correlation coefficient = 0.67, p-value < 1 × 10^−3^, **Figure 2H**). Such general co-regulation was further supported by the analysis of predicted TF binding motifs in both promoters (Pearson correlation coefficient = 0.63, p-value < 1 × 10^−3^) (**Supplementary Figure S10**). The profiles show groups of pcRNAs and coding genes as potential targets of the same general and developmental factors, suggesting that the coordinated expression between the pairs is largely due to the sharing of transcriptional regulatory elements.

The transcription factor occupancy analysis showed a striking enrichment of binding for CCCTC-binding factor (CTCF) in the regions adjacent to the TSSs of most pcRNAs (72% of pcRNA promoters contain a CTCF peak, **Figure 2H**,**3A**), and an enrichment with respect to other spliced lncRNAs and protein coding genes (**Figure 2H,3A**).

**Figure 3:**
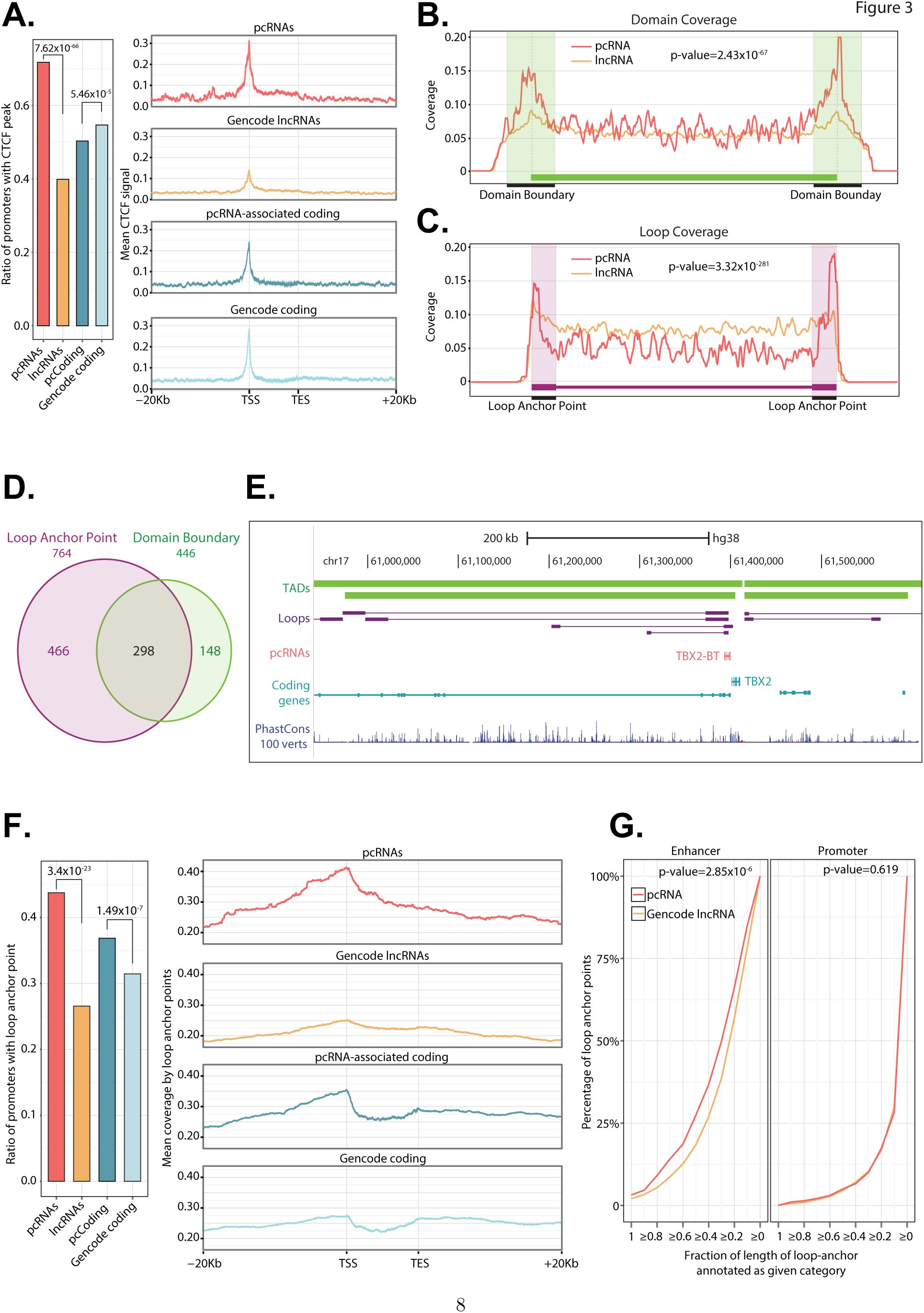
***A:** Bar chart showing the proportion of pcRNAs, pcRNA-associated coding genes, Gencode lncRNAs and Gencode coding genes with a CTCF peak (based on Encode ChIP-Seq data) overlapping their promoter. The p-values reported were calculated with hypergeometric tests. Right: CTCF peaks coverage of loci of pcRNAs, pcRNA-associated coding genes, Gencode lncRNAs and Gencode coding genes. The plots report the loci from 20kb upstream of the transcription start site (TSS) to 20kb downstream of the transcription end site (TES). For visualization purposes these profiles show the coverage of a random sample of 5000 Gencode lncRNAs and 5000 random Gencode coding genes. **B,C:** Aggregation density plots showing the distribution of the TSS of pcRNAs (red) and lncRNAs (orange) relative to chromatin topological domains (**B**) and chromatin loop anchor points (**C**). Domains and loop anchor points were defined based on HiC data. **D**: Venn diagram showing the number of pcRNAs whose promoter overlap a Loop Anchor Point (purple) or a Domain Boundary (green) **E:** Schematic representation of the TBX2 locus showing the pcRNA TBX2-BT and chromatin loops defined by HiC data (Rao et al., 2014). Modified from a screenshot of the UCSC genome browser. **F:** Bar chart showing the proportion of pcRNAs, pcRNA-associated coding genes, Gencode lncRNAs and Gencode coding genes with a HiC loop overlapping their promoter. The p-values reported were calculated with hypergeometric tests. Right: HiC loops coverage of loci of pcRNAs, pcRNA-associated coding genes, Gencode lncRNAs and Gencode coding genes. The plotted genomic regions encompass the loci from 20kb upstream of the TSS to 20kb downstream of the transcription end site (TES). For visualization purposes these profiles show the coverage of a random sample of 5000 Gencode lncRNAs and 5000 random Gencode coding genes. **G:** Cumulative distribution plot showing the percentage of distal genomic regions in contact with pcRNA promoters (y-axis) as a function of the fraction of length of loop-end annotated as Enhancer (left) or Promoter (right). For example, the “>0.4” point (x-axis) of the red line in the first plot indicates that ~37% (y-axis) of the distal genomic regions in contact with pcRNA promoters is annotated as Enhancer for 40% or more of their length. Promoters of pcRNAs are significantly more often in contact through loops with enhancer elements compared to generic Gencode lncRNAs (p-value 2.85x10^−6^). The indicated p-values were calculated using the Kolmogorov-Smirnov test*.

The genome is highly structured and gene expression influences and is influenced by the topological organization of the chromatin (Cavalli and Misteli, 2013). This organization is largely dictated by CTCF binding and includes individual chromatin loops and distinct regions with increased frequencies of genome looping contacts defined as topologically associating domains (TADs) (Rao et al., 2014). We thus interrogated high resolution, genome-wide topological maps (Rao et al., 2014) to establish the relative positioning of pcRNAs and their coding genes relative to genomic loops.

We found that pcRNAs are preferentially located at the boundaries of TADs and chromatin loop contact points (or “loop anchor points”) (**Figure 3B,C**) (Rao et al., 2014). In particular, we noticed that a remarkable proportion of pcRNAs (912 out of 1700 pcRNAs isoform TSSs, 54%) have a promoter that overlaps a TAD boundary (446 pcRNAs) and/or directly intersects a loop anchor point (764 pcRNAs, **Figure 3D**). For example, the pcRNA *TBX2-BT* and other pcRNAs associated with important developmental genes lie at TAD boundaries and overlap multiple loop anchor points (**Figure 3E**, **Supplementary Figure S11**).

Strikingly, the proportion of pcRNA promoters that overlap a TAD boundary or a loop anchor point is significantly higher than that of Gencode spliced lncRNAs (p-value=3.4 × 10^−23^, **Figure 3F**, **Supplementary Figure S12A,B**). Similarly, the promoters of pcRNA-associated coding genes are also significantly enriched in TAD boundaries or loop anchor points compared to other protein coding genes, although to a lesser extent than pcRNAs. Interestingly, we found that the location where loop anchor points are found around and within pcRNAs peaks distinctively at their TSS (**Figure 3F**), consistent with the observed positioning of CTCF binding sites and pcRNA TSS distribution within TADs and loops (**Figure 3A-C**).

Given this marked and precise association of their promoters with loop anchor points, we defined this group of 764 pcRNAs as ‘topological anchor point RNAs’ (tapRNAs), representing the subset of pcRNAs whose promoters overlap a loop anchor point (**Figure 3D**).

To gain further insights into the function of these promoter loops we analyzed the genomic regions they contact through looping. Based on the chromatin state segmentation by HMM from ENCODE/Broad for 9 cell lines (Ernst et al., 2011), we found that these loop anchor points are marked by multiple chromatin states and that a remarkable proportion appears to overlap marks of active transcription and/or enhancers (**Supplementary Figure S12C**). Interestingly, pcRNAs are significantly more likely to be in contact with enhancer elements through such loops, compared to Gencode lncRNAs (p-value=2.85 × 10^−6^; **Figure 3G**).

This is not the case for the contact with promoter elements, transcribed regions or other HMM defined genomic regions (**Figure 3G**, **Supplementary Figure S12D**). These results indicate that the tapRNAs that are at loop anchor points are likely to be associated with enhancer sequences present on the other end of the loop.

### 2.6 Conserved domains and motifs in tapRNAs

In order to establish any commonality of sequence that may provide clues to the function of tapRNAs, we explored their sequence conservation by direct RNA alignment. We found that 73% of tapRNAs show high conservation in patches of sequence between human and mouse (see Supplementary Methods), while the other 27% lack these highly conserved patches (**Figure 4A**). We divided the conserved 73% into (a) high-(b) medium-and (c) low-conservation-region tapRNAs and carried out a GO enrichment analysis on their associated coding genes. In all cases, the predominant category of genes associated with these tapRNAs was linked to development. In contrast, the 27% of tapRNAs with no highly conserved segments (d) were associated with genes showing no strong enrichment in functional categories. These data suggest that tapRNAs associated with developmental transcription factor genes have more significant sequence conservation than other tapRNAs. Indeed, tapRNAs as a class show more sequence conservation than lncRNAs in general (**Figure 4B**).

**Figure 4:**
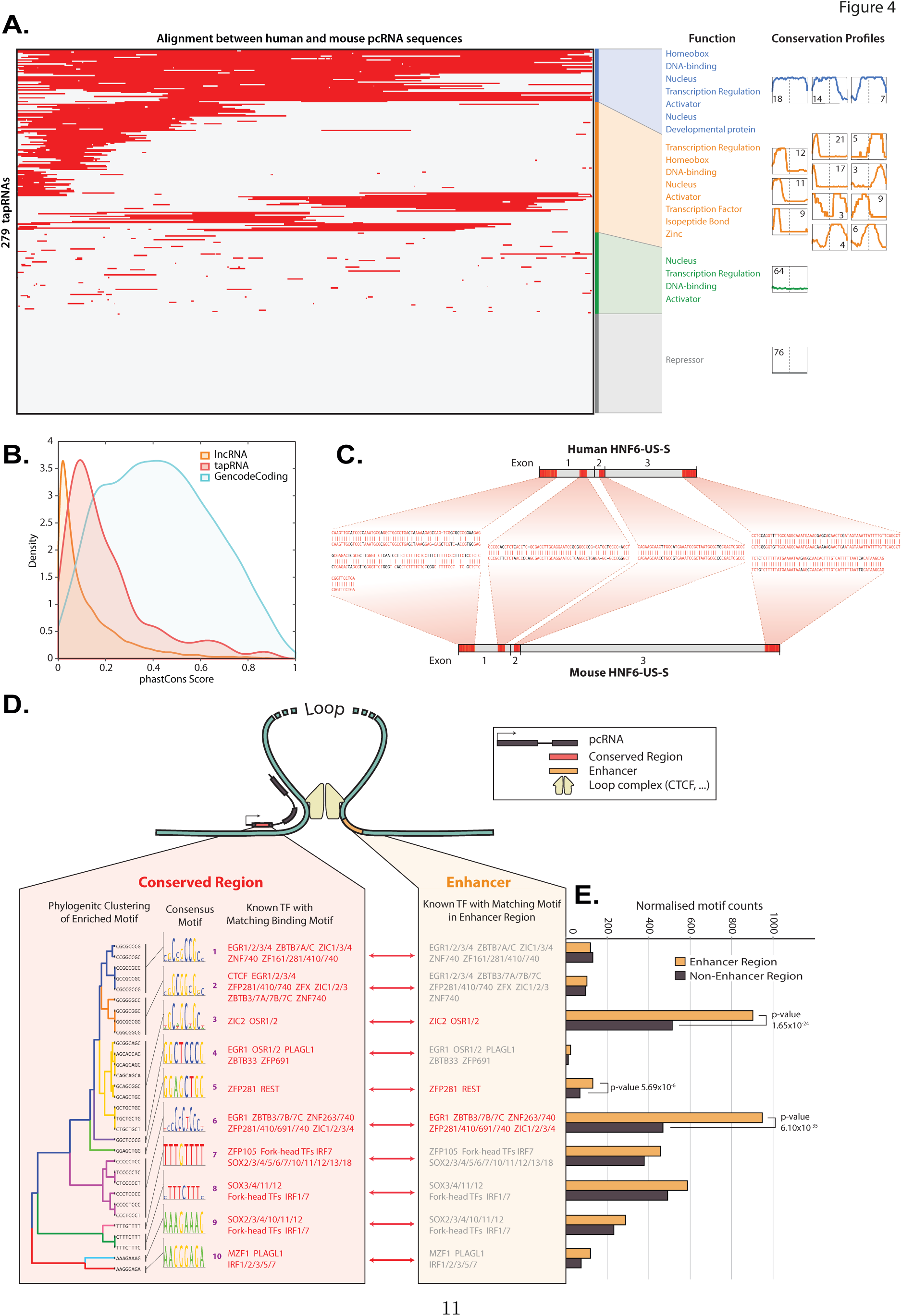
***A:** Clustered heatmap of conserved domains in transcribed tapRNAs. Aligned sequences (shown in red) in 279 non-redundant tapRNA isoforms are clustered (Euclidean distance). Sixteen minor clusters were identified and grouped into four major clusters. Each minor cluster’s centroids are shown with the number of tapRNAs belonging to each minor cluster. 39 tapRNAs (top group: blue) have more than 73% conserved domain in their transcribed sequences. Functional category annotation search finds that tapRNAs of top group are highly related to developmental proteins or Homeobox proteins. In contrast, 76 tapRNAs of the bottom cluster (grey) do not have any sequence conservation and do not show significant common functionality. There are also some minor groups in which position specific conservations are clearly present (e.g. 5’-end specific or 3’-end specific). **B:** Comparison of conservation between tapRNAs, lncRNAs and protein coding genes. The curves are Kernel Density Estimation (KDE) of conservation scores calculated from the phastCons multiple alignments of 100 vertebrate species. **C:** Example of conserved domains in a pcRNA. Identical sequence alignments (conserved domains) between human and mouse HNF6-US-S tapRNAs are represented in red, with RNA sequence alignments shown. **D:** Enriched TF-binding motif in both conserved domain of tapRNAs and enhancer region of loop anchor point. 32 significantly enriched 8-mer motifs (see **Supplementary Figure S13**; p-value 1 × 10^−4^) in conserved domains in tapRNAs are identified and clustered into 10 consensus motifs*. De novo *motif analysis discovers known TFs with matching binding consensus motifs. Seven out of ten consensus motifs are part of binding motifs of Zinc Finger proteins. The other three consensus motifs are part of binding motifs of developmental regulatory proteins. **E:** Extended motif search in enhancer regions of the other end of loop anchor points found significant enrichments of Zinc Finger protein motifs*.

The fact that we can detect conserved sequence domains within the tapRNA class (**Figure 4C**), prompted us to examine whether there are any sequence motifs in common between them. Motif enrichment analysis identified 32 8-mer motifs that were significantly more represented in the conserved domains of tapRNAs relative to non-conserved sequences (**Figure 4D**, **Supplementary Figure S13**). Closer inspection indicated that these 32 motifs were related and could be sub-categorized into 10 consensus motifs, which were reminiscent of transcription factor binding sites. All the 10 motifs have the potential to bind transcription factors, the predominant type being factors with Zinc Finger (ZF) domains (**Figure 4D**). Nevertheless, inspection of the available ChIP-seq data did not detect proteins of this category binding on the DNA corresponding to these regions in tapRNAs (data not shown). This raises the possibility that the motifs seen within the conserved domains of tapRNAs represent distinct RNA motifs.

Given that many tapRNA genes are at chromatin loop end points which have enhancers at the other end of the loop, we inspected whether the motifs enriched in conserved domains of tapRNAs are also enriched within the enhancer sequences found at the other end of the loop. We show (**Figure 4D,E**) that three out of the ten motifs enriched in tapRNAs are also enriched within enhancers on the other end of the loop anchor point. Strikingly, these three motifs are all motifs that have the potential to bind ZF proteins, including binding motifs for proteins associated with chromatin looping and enhancer function such as Znf143 and Zic2 (Heidari et al., 2014; Luo et al., 2015; Xie et al., 2013), which may be important for the function or regulation of these tapRNAs.

### 2.7 Functional analysis of positionally conserved RNAs

#### Co-regulation of gene expression

Given that pcR-NAs and their neighboring gene are co-expressed in a tissue-specific manner, we investigated their ability to regulate each other’s expression. First, we tested this hypothesis on a set of liver specific master regulators, *FOXA2* and *HNF1A*, and their associated pcRNAs. ChIP-sequencing data (Ballester et al., 2014; Down et al., 2011) shows that the promoters of the *FOXA2* gene and *FOXA2-DS-S* pcRNA are occupied by a similar set of transcription factors, and importantly, the same factors (*FOXA1*, *FOXA2*, *HNF4A* and *HNF6*) that are key regulators of liver differentiation (**Figure 5A**, **Supplementary Figure S14**). This again suggests that the mechanism of co-expression and co-induction is due, at least in part, to specific factors concomitantly regulating the expression of both the coding and non-coding transcripts.

**Figure 5:**
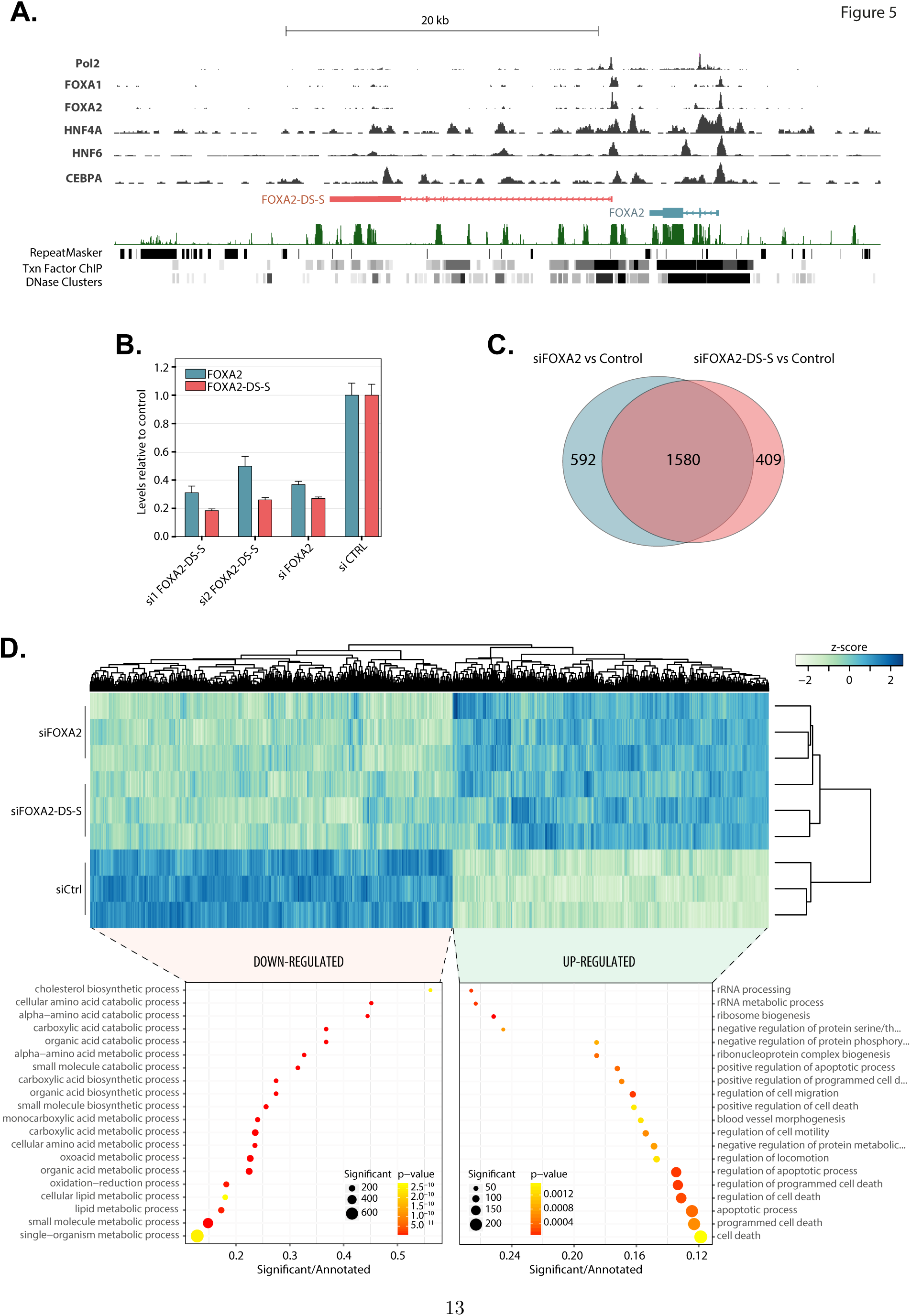
***A:** Screenshot from the Dalliance genome browser (Down et al., 2011) showing the FOXA2 locus with tracks displaying coverage data for ChIP-Seq experiments for Pol2, FOXA1, FOXA2, HNF4A, HNF6 and CEBPA. The ChIP-Seq tracks were produced by the ENCODE project on HepG2 cells. **B:** Real Time PCR data showing the expression of* FOXA2 *and* FOXA2-DS-S *in Huh7 cells upon knock-down. Si1-and si2-FOXA2-DS-S indicate two different, non-overlapping siRNAs designed against* FOXA2-DS-S*. The data is expressed relative to the expression of the control transfected with scrambled siRNAs; the error bars indicate the SEM across three replicate experiments. **C:** Venn diagram showing the number of significantly differentially expressed genes (adjusted p-value <0.05 and log2 fold change > or < 1.25) in the microarray experiment on Huh7 knock-down of* FOXA2 *or* FOXA-DS-S*. **D:** Heatmap showing microarray data upon knock-down of* FOXA2 *or* FOXA-DS-S *in Huh7 cells. The color-scale indicates normalized intensities (z-score). The heatmap contains all genes that were significantly altered (adjusted p<0.05) upon knock-down of either* FOXA2 *or* FOXA-DS-S*. The scatter plots in the lower part of the panel show GO enrichment data for genes that were significantly down (left) or up-regulated (right) in either siFOXA2 or siFOXA-DS-S*.

We next investigated whether the *FOXA2-DS-S* pcRNA can affect the expression of the associated coding gene, finding that *FOXA2-DS-S* is necessary for the full expression of the *FOXA2* gene (**Figure 5B**, **Supplementary Figure S15A**), since its down-regulation by RNA interference results in the reduction of *FOXA2* expression in Huh7 liver cells (**Figure 5B**), A549 lung cells (**Supplementary Figure S15A**) and HepG2 liver cells (data not shown). Interestingly, knock-down of *FOXA2* with a validated siRNA also leads to the down-regulation of *FOXA2-DS-S* (**Figure 5B**, **Supplementary Figure S15A**).

These results indicate that *FOXA2* not only auto-regulates by binding its own promoter (**Supplementary Figure S14**), but that it also affects the expression of the associated pcRNA. At the same time, the associated pcRNA is necessary for the expression of the *FOXA2* gene in different cell types, thus providing a positive feedback-loop and suggesting inter-dependence. This also raises the possibility that the pcRNA acts locally, *in cis*, in order to mediate its effect. Microarray analysis of the global transcriptional effects of *FOXA2-DS-S* or *FOXA2* knock-down showed a large overlap in the repertoire of affected genes (**Figure 5C,D**). This suggests that the major target for *FOXA2-DS-S* is the *FOXA2* gene, although there may be additional direct targets yet to be identified. These findings were recently independently supported by the cis-regulation of *FOXA2* in differentiating definitive endoderm cells by *FOXA2-DS-S* (termed in this study as *DEANR1*, or ‘definitive endoderm-associated lncRNA1’), indicating that this lncRNA regulates *FOXA2* in different endoderm-derived tissues (Jiang et al., 2015).

We also obtained similar results for two other tested pcRNAs in different cell lines (*POU3F3-BT* and *NR2F1-BT*) and also with a second liver factor *HNF1A* (**Supplementary Figures S6H** and **S15A-D**). In each of these cases, knock-down of the pcRNA reduces the expression of the associated coding gene. Interestingly, knock-down of *HNF1A-BT1* reduces the expression of *HNF1A* (**Supplementary Figures S6H**) but ectopic over-expression of full-length *HNF1A-BT* in human liver cells had no effect on the associated gene, even though very high levels of over-expression were achieved (data not shown), again suggesting a *cis*-based context-dependent mode of regulation.

#### Involvement in cancer

The Nanostring analysis of the cancer cell panel also demonstrated specific expression of pcRNAs, including many of the tapRNAs, and associated genes in different cancer lines (**Supplementary Figure S16A-E**). lncRNAs are emerging as important players in disease (Amaral et al., 2013; Balbin et al., 2015) and many genes with roles in development have been previously linked to cancer and other disorders (Dickerson et al., 2011). We therefore investigated the possible involvement of different pcRNAs in disease, specifically cancer. To further explore this association, we first performed a meta-analysis of the expression of pcRNAs in normal versus tumor samples in 63 microarray studies. After re-annotating the microarrays to identify probes targeting pcRNAs, we identified 203 pcRNAs significantly differentially expressed in tumors compared to normal tissues (**Figure 6A**, **Supplementary Table S6**) (see Supplementary Methods). These included known cases of lncRNAs involved in different cancers, such as *GAS5*, *DLEU2*, *PART1* and *MEG3* (Pickard and Williams, 2015).

**Figure 6:**
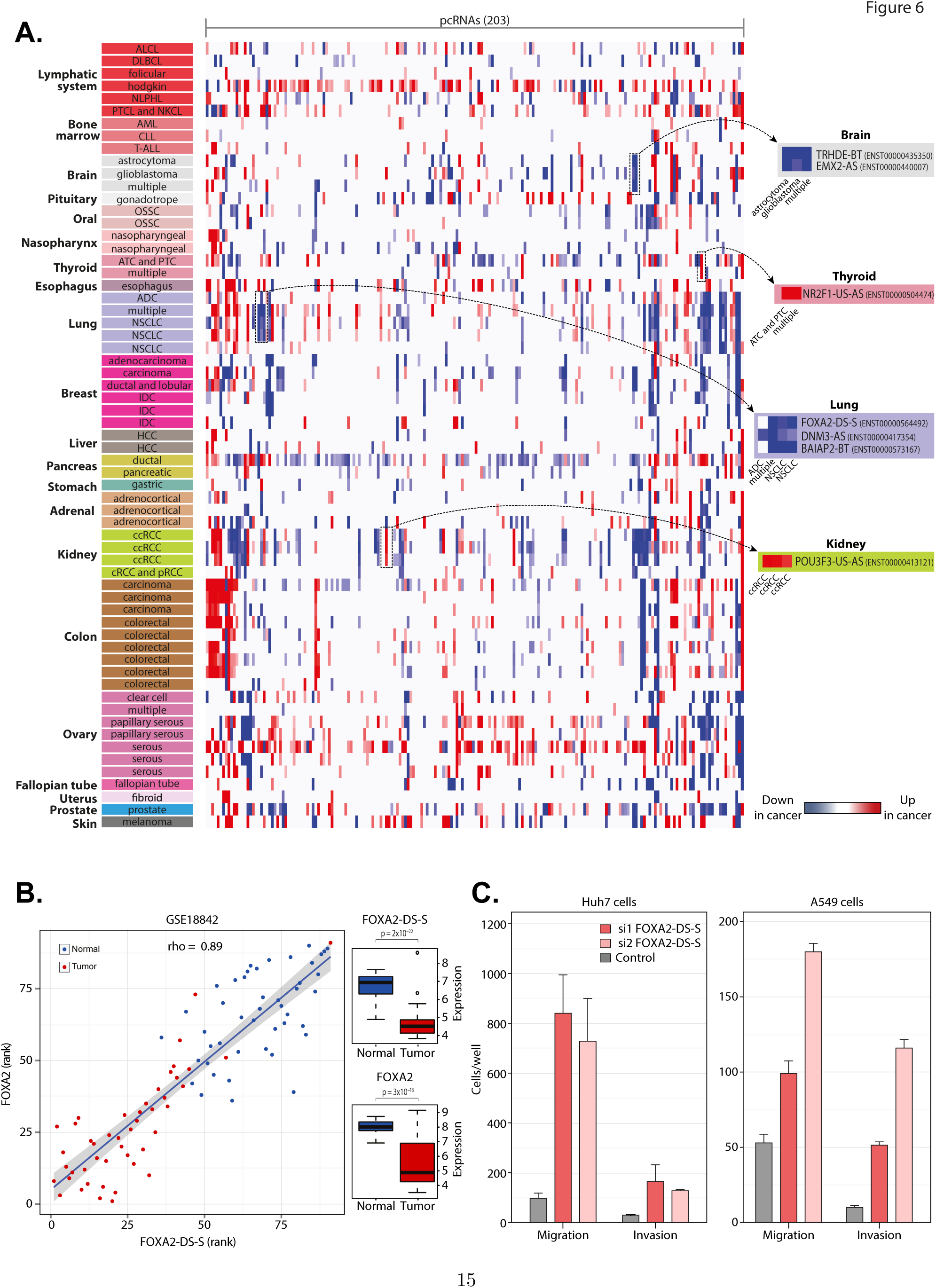
Figure 6: ***A:** Heatmap showing pcRNAs differentially expressed in cancer microarray studies. Student t-test (p-value < 0.005 and fold-change > 1.25) was used to identify pcRNAs (columns) that were up (red) or down-regulated (blue) in tumors compared to normal tissues (rows) (see Supplementary Table S6). Examples of pcRNAs associated with specific loci are shown. **B:** Spearman correlation between the expression of* FOXA2 *and* FOXA2-DS-S *in lung cancers (GSE18842 dataset). Tumor and normal individual samples are represented as blue and red dots, respectively. Boxplots on the right show that both transcripts are down-regulated in tumor compared to normal samples (Student’s t-test p-values are indicated). **C:** Invasion and migration assay analysis of Huh7 cells upon knock-down of* FOXA2-DS-S *using two different siRNAs (si1 and si2) compared to negative control siRNA*.

Close inspection of specific pcRNA-gene pairs with altered expression in primary tumors showed that a number of pcRNAs are consistently down-or up-regulated in cancer compared to normal tissues (**Supplementary Table S6**). For example, *FOXA2* and *FOXA2-DS-S* were found significantly down-regulated in lung tumor compared to normal samples (p-value 3 × 10^−16^ and 2 × 10^−22^ respectively), effectively separating tumor samples from controls (**Figure 6B**). Recent studies have correlated reduced *FOXA2* expression in hepatocellular carcinoma (HCC) with worse clinical outcome and identify this transcription factor as a tumor suppressor and inhibitor of epithelial-to-mesenchymal transition (EMT) in human lung cancers (Tang et al., 2011; Wang et al., 2014b). Here, our data also suggest the implication of the pcRNA *FOXA2-DS-S* in this process (**Figure 6B**).

To investigate whether the *FOXA2-DS-S* pcRNA has a potentially causative effect in cancer, we knocked-down *FOXA2-DS-S* in Huh7 and A5A9 cells (**Figure 6C**, **Supplementary Figure 15E**). A dramatic increase in both cell invasion and migration capacity was observed, supporting a tumor suppressor function for *FOXA2-DS-S* transcripts similar to the reported role of *FOXA2* in cancer cells (Basseres et al., 2012; Tang et al., 2011) (**Figure 6C**, **Supplementary Figure 15E**). These data further support the close functional link between the pcRNA and its associated gene. The fact that the *FOXA2-DS-S* pcRNA has the same effect on the phenotypic characteristics of cancer cells as its associated gene, is consistent with its positive effect on the expression of *FOXA2* and the observation that it regulates a similar cohort of genes (**Figure 5C,D**).

Finally, we also investigated the role of two other pcRNA-gene pairs in invasion and migration characteristics of cancer cells. We found that knock-down of *NR2F1-BT* and *NR2F1* or *POU3F3-BT* and *POU3F3* similarly reduced the invasion/migration potential of U251MG glioblastoma and U2OS osteosarcoma cells (**Supplementary Figure S15F-H**). Thus, in the case of pcRNAs *NR2F1-BT* and *POU3F3-BT*, their effect on cell invasion/migration suggests that they, like their associated gene, may have oncogenic roles in these cells. Taken together these data highlight the fact that certain pcRNAs can be considered as potential targets for cancer and that their expression pattern may act as a marker for the disease.

## 3 Discussion

### 3.1 Positional conservation identifies lncRNAs with common characteristics linked to developmental genes

In this work, we systematically identify, on a genome-wide scale, lncRNAs with conserved promoters in mouse and human that are positionally conserved in respect to neighboring coding genes. We catalogue 665 lncRNA promoters that are linked to 626 coding genes and define these as positionally conserved. This analysis has identified a subset of lncRNAs with a close relationship to a very specific cohort of neighboring protein-coding genes, predominantly comprised of transcription factors with well-established roles in development.

Our analysis of positionally conserved lncRNAs and their associated developmental transcription factors has identified common characteristics within this group, which indicate that they are (a) expressed in the same restricted tissues in mouse and human and (b) co-induced when cells are stimulated with a differentiation signal, (c) have promoters bound by similar transcription factors, (d) can affect each others expression, (e) affect invasion and migration of cancer cells in similar ways, and (f) are part of a new subclass of lncRNAs, which are positioned at topological anchor points (tapRNAs) and have a specific set of motifs enriched within their conserved sequence.

Our dataset encompasses hundreds of previously uncharacterized positionally conserved RNAs and includes a few lncRNAs that have been studied by others on a case-by-case basis (e.g. *HOTTIP* (*HOXA13-AS*), *HOTAIRM1* (*HOXA1-AS*), *NKX2-1-AS and FOXA2* (*DEANR1*, *FOXA2-DS-S*)) (Balbin et al., 2015; Deng et al., 2016; Jiang et al., 2015; Wang et al., 2011; Zhang et al., 2009). In all these studies the pcRNAs were shown to regulate developmental genes in different biological contexts, in a manner implying local regulation. Our investigation of positionally conserved RNAs, as well as recent analysis of syntenic RNAs defined by others (Amaral et al., 2009; Dinger et al., 2008; Hezroni et al., 2015), highlights a common grouping of lncRNAs that have close functional connections with developmental transcription factors. Given this spatial and functional connection, in this work we have annotated pcRNAs using a systematic nomenclature that reflects the orientation of the pcRNA relative to the associated coding gene. We believe that this catalogue will be a valuable resource in the individual characterization of these important coding genes and their associated non-coding RNA genes.

### 3.2 Co-regulation of positionally conserved RNAs and developmental genes

Our analysis shows that pcRNAs and their associated coding genes are co-expressed in a tissue specific manner and co-induced after treatment with differentiating agents, consistent with previous observations by us and others that show co-regulation of lncRNAs and neighboring genes in different species (e.g. (Cabili et al., 2011; Dinger et al., 2008; Goff et al., 2015; Hezroni et al., 2015; Mercer et al., 2008)). The fact that these lncRNAs and associated coding genes have similar transcription factor binding profiles within their promoters (**Figure 2H** and **Supplementary Figure S10**), and that these transcription factors can be tissue specific (**Figure 5A**, **Supplementary Figure S13**), provide some insight into the underlying mechanism. As an example we have analyzed the liver specific gene *FOXA2* and its pcRNA *FOXA2-DS-S*. The promoters of each of these are bound by liver specific factors, such as HNF4A, HNF6 and FOXA2 itself. Consistent with this, we find that the transcription factor *FOXA2* can regulate both its own transcription and that of the *FOXA2-DS-S* pcRNA. This example indicates a mechanism by which one tissue specific factor (*FOXA2*) can regulate both its own coding gene and the associated pcRNA in a positive feedback loop to exert co-expression in liver cell lines.

We have investigated the regulation of *FOXA2* by the *FOXA2-DS-S* pcRNA, and we see a positive effect on its expression levels. This positive regulatory effect could be extended to other pcRNAs and different cell types, in line with previous reports (e.g. (Jiang et al., 2015) and references above). These results do not exclude the possibility that pcRNAs may also have an effect on other neighboring or distal genes (*in trans*), or that the regulation is repressive for gene expression, as is the case for imprinted lncRNAs and other transcripts (Goff et al., 2015; Guil and Esteller, 2012; Kanduri, 2015). In addition, previous studies have shown that genetic removal of lncRNA loci only in some cases affect the expression of neighboring genes in specific developmental stages *in vivo*, suggesting that these loci are not required for cis-regulation, although mechanisms such as genetic compensation and robustness may be involved (Rossi et al., 2015). Nevertheless, the functional evidence presented and the numerous cases of pcRNA positive correlation of expression with the associated gene in both mouse and human support the hypothesis that positive and local regulation is an important mode of pcRNA regulation.

### 3.3 Positionally conserved RNAs and cancer

Many developmental genes are implicated in cancer, either positively or negatively. The expression analysis of pcRNAs in primary cancers indicates that they are over-or under-expressed in particular types of cancer, consistent with other studies (Balbin et al., 2015; Iyer et al., 2015). Importantly, we show here that manipulation of the levels of a pcRNA has an effect on the metastatic characteristics of cancer cells *in vitro*. These phenotypic effects can be either positive or negative, but in each tested case the associated coding gene and the pcRNA have a similar effect on the metastatic characteristic of cancer cells. Collectively, these data suggest that pcRNAs have the potential to be causally linked with the disease, and that the coding gene they are associated with is a key target. We cannot exclude the possibility that other targets of different pcRNA are drivers of the cancers. However, pcRNAs or the pathways that regulate them may represent targets for therapeutic intervention given their highly specific expression and alteration in particular cancers.

### 3.4 Topological anchor point RNAs (tapRNAs)

The most striking feature of positionally conserved RNAs is their enrichment in tapRNAs, defined as RNAs found at chromatin loop anchor points. These architectural landmarks are commonly occupied by the CTCF boundary factor (Rao et al., 2014) and indeed we find an enrichment of CTCF binding in the promoters of pcRNAs.

The presence of genes within chromatin loops has already been proposed as an indication of increased and coordinated expression and, although many such loops do not show tissue specificity, other topology-dependent events or finer-scale topological contacts likely regulate tissue-specific and fine-tuned expression (Dowen et al., 2014; Lonfat et al., 2014; Rao et al., 2014). LncRNAs are rapidly emerging as important targets and actors in the regulation of genome topology and nuclear architecture (Rinn and Guttman, 2014). These include lncRNAs whose functions are regulated by CTCF and chromatin looping structures (e.g. (Engreitz et al., 2013; Maass et al., 2012; Sopher et al., 2011; Spencer et al., 2011; Yang et al., 2016)), but also well-characterized pcRNAs such as *XIST* and an increasing number of more recently identified cis-and trans-acting lncRNAs such as *FIRRE*, *TSIX*, *KCNQ1OT1*, *CCAT1-L*, *RUNXOR*, *IRAIN*, ncRNA-a, *HoxBlinc* and *HOTTIP*, which have been shown to influence topological domains and chromosome conformation, for example by modulating the binding of transcription factors, Mediator and CTCF complexes (Deng et al., 2016; Hacisuleyman et al., 2014; Kung et al., 2015; Lai et al., 2013; Minajigi et al., 2015; Sun et al., 2014; Wang et al., 2014a; Wang et al., 2011; Xiang et al., 2014; Yang et al., 2015; Zhang et al., 2014).

The finding that over 50% of positionally conserved RNAs in our dataset are tapRNAs suggests that these promoters and transcripts are more generally connected to the structure and function of chromatin architectural domains. This is especially relevant given the robust conservation of chromatin conformation in syntenic regions, in which conserved CTCF-Cohesin binding sites are enriched at the borders of topological domains (Vietri Rudan et al., 2015). In addition, important developmental genes are positioned and regulated at such boundaries (Fabre et al., 2015) and these structures can be especially implicated in developmental diseases and cancer (Hnisz et al., 2016; Ibn-Salem et al., 2014; Katainen et al., 2015; Lupianez et al., 2015; Maass et al., 2012). The mechanisms may involve the conserved tapRNA promoters present at the boundaries, as many of these harbor highly conserved sequences that potentially act as cis-regulatory elements. These are exemplified by the promoters of *SOX2OT* and other pcRNAs associated with developmental genes, such as *Dlx1as* and *Evf1/2* (Amaral et al., 2009; Bond et al., 2009; Dinger et al., 2008; Feng et al., 2006), although conserved promoters and lncRNA loci may exhibit both DNA-and RNA-dependent regulatory functions (Bond et al., 2009). TapRNAs themselves could be either influencing the formation of a topological anchor point or be expressed at that precise location in order to affect a local function in a cell-type specific manner (although these alternatives are not mutually exclusive).

Moreover, our observation that enhancers are often found on the other end of the loop containing a tapRNA may be relevant to the mechanism of action of these transcripts. Looped chromatin structures form discrete chromatin domains and regulatory “neighborhoods” that influence the local regulation of developmental and cell-identity genes, tissue-specific enhancers and other regulatory elements (Dowen et al., 2014; Narendra et al., 2015). Since CTCF brings together contact points on both side of a loop, the tapRNA can be in close proximity to enhancers and thereby contribute to enhancer regulation in a number of ways. TapRNAs may directly deliver or remove proteins necessary for looping and enhancer function; and interact with DNA elements at the enhancer or with eRNAs transcribed from these (Arner et al., 2015; Kim et al., 2015; Pnueli et al., 2015; Sigova et al., 2015). These different potential mechanisms of action warrant experimental validation, and may be mediated by specific sequence motifs within tapRNAs, as discussed below.

### 3.5 Conserved domains and motifs within tapRNAs

Analysis of sequence conservation among tapRNAs has shown that this subclass of RNAs is more conserved than the bulk lncRNAs. Within the conserved sequences of tapRNAs, ten degenerate motifs were particularly enriched. These are similar to motifs for the binding of transcription factors, predominantly of the C2H2 Zinc Finger (ZF) family. Interestingly, ZF domains of the C2H2 family have the ability to bind both DNA and RNA (Brown, 2005). Thus these motifs may represent part of the common functional characteristics of tapRNAs, which could involve interaction with ZF proteins or alternatively, base pairing with homologous RNA or DNA sequences.

We find that ZF motifs within tapRNAs are also enriched at enhancer sequences at the other end of the loop. For example, one of these enhancer DNA motifs (for the ZF protein ZIC2) has been shown to act as a repressor of enhancer activity in a developmental context (Luo et al., 2015). This raises the possibility that the mechanism of action of ZF motifs within tapRNAs is related to the presence of similar ZF motifs and DNA bound proteins at enhancers and on chromatin topology. In this way, tapRNA ZF motifs could include the sequestration or delivery of ZF factors to enhancers, for example involving the formation of RNA-DNA hybrids or triplex structures (Mondal et al., 2015; O’Leary et al., 2015). In all these scenarios, the tapRNAs could have a positive effect on the transcription of the associated developmental coding gene, depending on whether the ZF transcription factor is an activator or repressor of transcription. Binding of the YY1 transcription factor to RNA has recently been demonstrated to play a role at enhancers (Sigova et al., 2015), supporting the argument that RNA-transcription factor interactions may influence enhancer activity.

### 3.6 The “extended gene” concept in regulatory evolution

Positionally conserved RNAs defined here include a number of well-characterized lncRNAs (such as *XIST*, *H19*, *AIRN*, and *MEG3*), as well as more recently characterized syntenic RNAs (*HOXA13-AS*, *HOXA1-AS*, *EVF2*/*DLX6OS*, *NKX2-1-AS and DEANR1/FOXA2-DS-S*) linked to genes with important developmental roles. These examples and the data presented here conform to a principle where coding and noncoding genes and associated regulatory sequences have evolved in a manner that favors genomic association and co-expression, and in which the noncoding RNA can have a role in the regulation of the coding gene (and sometimes vice-versa) and related developmental pathways.

The reason for such apparent co-evolution may be that the more evolutionary plastic lncRNAs and associated cis-regulatory elements – with different evolutionary constrains compared to protein coding sequences - may act as substrates for the alteration of regulatory functions, which might provide advantages in determining a new developmental direction. Thus, positional conservation may define examples of an “extended gene” paradigm, where a lncRNA gene and its associated regulatory elements are linked in genomic position and function to a coding gene. We believe there may be many more instances of lncRNA loci involved in an extended gene structure besides the ones outlined in this work, as our analysis is inherently limited to a particular subset of syntenic spliced lncRNAs with conserved promoters and excludes numerous lncRNAs that do not meet our inclusion criteria, for example single exon or more clade-restricted transcripts and promoters.

Regardless of these limitations, our current analysis shows that only a small fraction (˜3%) of protein-coding genes have positionally conserved RNAs and that tapRNAs represent a large proportion of these. The fact that positional conservation identifies RNAs at topological anchor points, strongly suggests that the conservation is reflective of chromatin architectural features or mechanisms to regulate them. This type of architectural control may be particularly important for the expression of developmental genes and its regulatory dynamics across evolution.

## 4 Methods

### 4.1 pcRNA cloning

Cloning of human full-length HNF1A-BT1 transcript was performed using Gateway Technology (Thermo Fisher Scientific, cat. no. 12535-019) according to the manufacturer’s instructions. Briefly, 2μg total RNA from HepG2 cells was reverse transcribed in 20μl reaction using Superscript III Reverse Transcriptase (Invitrogen; cat. no. 18080044). Touchdown-PCR was performed using 2μl of the cDNA, mixed with 38.75μl water, 1μl of each primer (10μM) (Supplementary Table S7), 1.25μl dNTP (10mM), 5μl 10x Pfu Ultra reaction buffer and 1μl Pfu Polymerase (Stratagene; cat. no. 600380). PCR was performed in the following conditions: (i) 98°C for 30s, (ii) 98°C for 10s, (iii) 70-50°C for 30s, (iv) 72°C for 3min (with 20 cycles repeating steps (ii) - (iv), thereby decreasing the temperature of step (iii) 1°C per cycle); followed by (v) 98°C for 10s, (vi) 50°C for 30s, (vii) 72°C for 2min, (viii) 72°C for 5min, with 15 cycles repeating steps (v)-(vii). Nested PCR was performed with PCR products after gel extraction and 1:100 dilution. The cycling conditions were: (i) 98°C for 30s, (ii) 98°C for 10s, (iii) 59°C for 30s, (iv) 72°C for 2min, (v) 72°C for 5 min, with 30 cycles repeating steps (ii)-(iv). Gel purified PCR products were quantified and transferred into the pDONR221 entry vector (Life Technologies, cat. no. 12536-017). Transformations were performed according to the Gateway clonase protocol using *Escherichia coli* DH5α and the plasmids used in an LR reaction to generate the expression vectors using the LINC-EXPRESS plasmid (modified pLENTI6.3/TO/V5-DEST, kindly provided by John Rinn (Hacisuleyman et al., 2014).

### 4.2 Cloning of shRNAs

Short hairpin (sh)RNA design was performed using the Broad Institute RNAi Consortium software (www.broadinstitute.org/rnai/public/seq/search), which was used to select three or four different candidate shRNAs per pcRNA. We tested their knockdown efficiencies in transient transfections and the effective shRNAs as well as negative control were used in subsequent experiments (Supplementary Table S7). Cloning of the annealed shRNA oligos into pLKO vectors for shRNA Constructs was performed as described in the TRC Laboratory Protocol (Version 2/12/2013). Briefly, HPLC-purified oligomers were annealed using final concentrations of 3μM each in 1xNEB buffer 2 and used in ligations into pLKO vectors digested with AgeI and EcoRI (pLKO.1 puro (addgene #8453; (Stewart et al., 2003)) and Tet-pLKO-puro (addgene #21915; (Wiederschain et al., 2009)). Ligation products were used for transformations of *E. coli* DH5α and plasmids were purified and sequenced.

### 4.3 Cell culture

Cell lines (Supplementary Table S7) were acquired from ATCC and cultured at 37°C and 5% CO_2_ in the recommended media unless specified. Human hepatocellular carcinoma cells Huh-7 and HepG2 cells were grown in growth medium (Dulbecco’s modified Eagle’s medium (DMEM), 10% fetal bovine serum (FBS), 2 mM L-glutamine, 50 ug/ml penicillin and 50 ug/ml streptomycin) at 37°C and 5% CO_2_. Lung adenocarcinoma, Osteosarcoma and glioblastoma cells A549, U2OS and U251MG were cultured in DMEM supplemented with 10% FBS. Mouse embryonic stem cells (E14) were cultured in 1% gelatin-coted dishes in DMEM media with 2mM glutamine, 1mM sodium pyruvate, 1X non-essential amino acids, 10% FBS, 0.1 mM 2-mercaptoethanol, and supplemented with 1,000 U/mL LIF (ESGRO). Human teratocarcinoma NT2/D1 cells were cultured in DMEM supplemented with 10% FBS.

For induction NT2 cells and mESCs in media without LIF were treated with 10μM all-trans retinoic acid (Sigma R 2625) and harvested at different time points.

### 4.4 Knock-down

SiRNA oligonucleotides were ordered from Thermo Fisher Scientific (Supplementary Table S7). For over-expression of coding genes, plasmids were ordered from Origene (Supplementary Table S7). Transfections for siRNA-mediated knock-down experiments were performed according to the Lipofectamine RNAiMAX (Thermo Fisher Scientific, cat. no. 13778150) procedure. Briefly, the day before transfection 1 × 10^5^ cells were seeded in 2.5ml DMEM/10% FBS in 6-well plates. For each well, 50nM siRNA duplexes were diluted in 250μl Opti-MEM. 5μl Lipofectamine RNAiMAX were added to 245μl Opti-MEM and combined with the siRNA mix. After incubation for 10-20min at room temperature (RT) the mix was added drop-wise to the cells. Cells were incubated for 48h until harvest. For over-expression, the day before transfection 1 × 10^5^ cells were plated per 6-well in 2ml growth medium without antibiotics. Transfections were performed following the Fugene6 Transfection Reagent Protocol (Promega, cat. no. E2691). Briefly, 185μl medium were pre-mixed with 5μl transfection reagent per well in a 6-well plate. After 5min incubation at RT, 10μl plasmid DNA (1μg) were added to the mix and incubated for 30min at RT. Subsequently the transfection reagent/DNA mixture was added dropwise to each well. The transfected cells were incubated for 48h before processing.

### 4.5 RNA isolation and RT-PCR expression analysis

RNA from cell cultures was purified using Qiazol (Invitrogen) and RNeasy Mini Kit (Qiagen) or Direct-zol RNA MiniPrep kit (Zymo Research, cat. no. R2072) and treated with DNase I (Invitrogen), according to manufacturers’ instructions. Purified tissue RNA was purchased from Ambion (FirstChoice Human Total RNA Survey Panel, cat. no. AM6000) and Clontech (Mouse Total RNA Master Panel, cat. no. 636644) (Supplementary Table S7). RNA quality assessed using the 2100 Bioanalyzer (Agilent Technologies) prior to further use. cDNA preparation and quantitative real-time PCR (qPCR) analysis were performed as previously described (Dinger et al., 2008). Each experiment was performed in at least two biological replicates. Primers were designed spanning splice sites in most cases and for normalization of transcript expression levels, *B2M*, *ALAS1* or *GAPDH* primers were used (Supplementary Table S7).

### 4.6 Invasion-migration assays

2.5 × 10^4^ cells were plated in serum-free media in insert plate upper chamber with either non-or Matrigel-coated membranes (24-well insert; pore size, 8 μM; BD Biosciences cat. no. 354578 and 354480) for transwell migration and invasion assay, respectively. The bottom chamber contained DMEM with 10% FBS. After 24h, the bottom of the chamber insert was fixed and stained with crystal violet and cells on the stained membrane were counted under a microscope. Each membrane was divided into four quadrants, and an average from all four quadrants was calculated. Each assay was performed in biological triplicates.

## 5 Accession numbers

The microarray data have been submitted to ArrayExpress under the accession number: E-MTAB-4517.

## 6 Acknowledgements

This work was funded by programme grants from Cancer Research UK (C6/A18796) and European Research Council CRIPTON Grant (268569), and supported by a University of Cambridge and FAPESP grant (2014/50308-4) and Institutional funding by a Wellcome Trust Core Grant (092096) and Cancer Research UK Grant (C6946/A14492). We thank John Rinn for many constructive discussions and for providing the lncRNA over-expression vector.

